# Pharmacogenetic testing through the direct-to-consumer genetic testing company 23andMe

**DOI:** 10.1101/098541

**Authors:** Mengfei Lu, Cathryn M. Lewis, Matthew Traylor

**Affiliations:** Department of Medical and Molecular Genetics, King’s College London, 8^th^ Floor Tower Wing, Guy’s Hospital, Great Maze Pond, London, SE1 9RT, UK; SGDP Centre, Institute of Psychiatry, Psychology & Neuroscience, King’s College London, De Crespigny Park, Denmark Hill, London, SE5 8EF, UK

## Abstract

**Background:** Rapid advances in scientific research have led to an increase in public awareness of genetic testing and pharmacogenetics. Direct-to-consumer (DTC) genetic testing companies, such as 23andMe, allow consumers to access their genetic information directly through an online service without the involvement of healthcare professionals. Here, we evaluate the clinical relevance of such pharmacogenetic tests reported by 23andMe.

**Methods:** The research papers listed under each 23andMe report were evaluated, extracting information on effect size, sample size and ethnicity. A wider literature search was performed to provide a fuller assessment of the pharmacogenetic test and variants were matched to FDA recommendations. Additional evidence from CPIC guidelines, PharmGKB, and Dutch Pharmacogenetics Working Group was reviewed to determine current clinical practice. The value of the tests across ethnic groups was determined, including information on linkage disequilibrium between the tested SNP and causal pharmacogenetic variant, where relevant.

**Results:** 23andMe offers 12 pharmacogenetic tests, some of which are in standard clinical practice, and others which are less widely applied. The clinical validity and clinical utility varies extensively between tests. The variants tested are likely to have different degrees of sensitivity due to different risk allele frequencies and linkage disequilibrium patterns across populations. The clinical relevance depends on the ethnicity of the individual and variability of pharmacogenetic markers. Further research is required to determine causal variants and provide more complete assessment of drug response and side effects.

**Conclusion:** 23andMe reports provide some useful pharmacogenetics information, mirroring clinical tests that are in standard use. Other tests are unspecific, providing limited guidance and may not be useful for patients without professional interpretation. Nevertheless, DTC companies like 23andMe act as a powerful intermediate step to integrate pharmacogenetic testing into clinical practice.

## Introduction

Recent advances in technology have enabled comprehensive characterization of the genetic component underlying many complex diseases, traits, and responses to medication. This new information has enabled genetic testing to become more widely available in healthcare, and can be used to assess risk of inherited conditions and predict response to medication. Such testing has the potential to reduce drug-related adverse events, as well as to increase the effectiveness of drugs by assessing how sensitive an individual might be to a given pharmaceutical. Several pharmacogenetic tests have become standard clinical practice and others are supported by strong research evidence. However, challenges exist in moving pharmacogenetic testing from a research endeavor to point-of-care implementation.

Traditionally, genetic tests have been ordered in clinical settings but direct-to-consumer (DTC) genetic testing companies allow consumers to access their genetic information through an online service without the intermediary of a healthcare professional. A 2012 review of online companies offering pharmacogenetic testing identified eight companies providing at least one such test, either DTC or through a healthcare practitioner. 23andMe was the most comprehensive DTC pharmacogenetic testing company in 2012, and remains such today, although its portfolio of tests has reduced substantially since scrutiny by the FDA in 2013.

Pharmacogenetic testing is an interesting market for DTC companies, since test results only become actionable when a relevant drug is prescribed, which may be at the point of testing or many years later. This contrasts with genetic testing for disease variants where the increased – or decreased – risk is of immediate relevance for the participant. It may be cost-effective for individuals to undergo pharmacogenetic testing once in their adult life, and store the results, informing physicians whenever a new drug is prescribed so any relevant gene-drug associations may be checked. Such pre-emptive pharmacogenetic testing may become routine practice, with data stored in the electronic health record (EHR), but there are few current implementations of this.[1] The introduction of DTC genetic testing has raised significant controversy amongst clinicians, academic researchers and the general public.[2] One of the major concerns is the potential for misunderstanding and misinterpretation of the test results, particularly when pre- or post-test genetic counselling is not provided.

Here we study the pharmacogenetic tests offered by 23andMe to UK customers. 23andMe began offering health-related tests to a UK market in December 2014, when their US tests covered only ancestry testing, following FDA concerns over their health tests. Currently, 23andMe provides reports on over 100 health conditions and traits for UK customers. These reports include (1) recessive inherited variants for conditions such as cystic fibrosis and beta-thalassaemia, (2) dominantly inherited genetic risk factors like variants in the *BRCA1* and *BRCA2* genes, (3) drug response tests like abacavir hypersensitivity and warfarin sensitivity and (3) prediction of traits such as hair colour and earwax type. 23andMe genotypes consumer’s DNA on the Illumina HumanOmniExpress-24 format chip, with more than 715,000 single nucleotide polymorphisms (SNPs), together with custom content specific for their tests (https://www.23andme.com/en-gb/health/; accessed 3^rd^ November, 2016). Consumer reports list each relevant genotype together with a summary, and further information on the test. The criteria for including a genetic test in the 23andMe portfolio are that the drug response tests are *eligible* if there are either existing clinical practice guidelines provided by CPIC and other clinical organisations, or if information from regulatory agencies or in drug labels “acknowledges the impact of the genetic marker on drug response”. The pharmacogenetic tests are *potentially eligible* if 23andMe is able to recognise benefits of the results, and if at least three scientific research papers identify consistent clinical effects of the marker tested.[3]

23andMe currently provides reports on 12 pharmacogenetic tests for UK customers. Here, we evaluate the extent to which these tests 1) represent the latest scientific and clinical literature, 2) reflect recommendations from the U.S. FDA and CPIC, and 3) are applicable across different ethnic groups.

## Methods

### Data sources and search strategy

Each test within the Drug Response section of 23andMe UK genetic testing report was reviewed *(https://www.23andme.com/en-gb/health/reports).* For all 12 pharmacogenetic tests, the 23andMe website provides details of 1) the drug, disorder and variant(s) relevant for the pharmacogenetic test, 2) possible outcomes by genotype, 3) interpretation of test results, and 4) academic papers reporting the pharmacogenetic association. Full technical information on the SNPs used to determine test results are given, enabling each test to be assessed fully.

The research papers listed under each 23andMe report were evaluated, extracting information on effect size, sample size and ethnicity. A wider literature search was performed using PubMed and EMBASE assess the pharmacogenetic test fully. The variants reported by 23andMe were matched to the FDA Table of Pharmacogenomic Biomarkers in Drug Labelling.[4] Additional evidence from CPIC guidelines,[5] PharmGKB,[6] and Dutch Pharmacogenetics Working Group (DPWG)[7] was reviewed to determine current clinical practice. SNP allele frequencies and linkage disequilbrium (LD) between SNPs were determined using the 1000 genomes project data through SNAP (http://archive.broadinstitute.org/mpg/snap/) and LDlink (https://analysistools.nci.nih.gov/LDlink/).[8–10]

## Results

### Assessment of 23andMe Pharmacogenetic Tests

23andMe provides pharmacogenetic tests for responses to 12 drugs, which are listed in Table 1, together with the specific variants tested, the implicated gene, the phenotype tested for (adverse drug reaction (ADR), efficacy, dosage adjustment), and the FDA recommendation. The tests are from diverse clinical areas and drugs. Five tests are for variants in the cytochrome P450 genes *CYP2C19* and *CYP2C9*, which affect both drug efficacy and adverse events. Genotypes are classified by metaboliser level and CPIC guidelines for dose adjustment or recommendations for a different drug are given. The remaining seven tests are for drug toxicity and adverse events, such as abacavir hypersensitivity reactions which are associated with HLA-B*57:01. All tests are for variants in a single gene, except for warfarin tests which combine genotypes from three SNPs in *CYP2C9* and *VKORC1*. The test for acetaldehyde toxicity, leading to increased risk of oesophageal cancer, is a curious entry in the list, since the toxicity arises from a breakdown of alcohol, a recreational drug, and not a pharmaceutical compound as in the other 11 tests.

**Table 1:** Pharmacogenetic tests provided by 23andMe

| Drugs | Gene | Variant/Marker | SNP | Purpose of test | FDA/PharmGKB Guidance<sup>a</sup> |
| --- | --- | --- | --- | --- | --- |
| Abacavir | HLA-B | *57:01 | rs2395029 | Hypersensitivity reactions | Genetic testing required |
| Acetaldehyde | ALDH2 | *2 | rs671 | Toxicity and risk of oesophageal cancer | No recommendation |
| Clopidogrel (Plavix®) | CYP2C19 | *2<br>*3<br>*4<br>*8<br>*17 | rs4244285<br>rs4986893<br>rs28399504<br>rs41291556<br>rs12248560 | Efficacy and ADRs | Genetic testing recommendation |
| Fluorouracil (Adrucil®) | DPYD | *2A | rs3918290 | Toxicity and ADRs | Actionable PGx |
| Peginterferon alpha (PEG-IFN-alpha) & ribavirin (RBV) | 19q13 region |  | rs8099917 | Hepatitis C treatment failure | Actionable PGx |
| Phenytoin | CYP2C9 | *2<br>*3 | rs1799853<br>rs1057910 | Sensitivity and Dosage adjustment | Actionable PGx |
| Proton Pump Inhibitor (PPI) | CYP2C19 | *2<br>*3<br>*4<br>*8<br>*17 | rs4244285<br>rs4986893<br>rs28399504<br>rs41291556<br>rs12248560 | Efficacy and Dosage adjustment | Informative PGx |
| Pseudocholinesterase Deficiency | BCHE | F1<br>F2<br>A | rs28933389<br>rs28933390<br>rs1799807 | ADR - extended paralysis and apnoea | Actionable PGx |
| Simvastatin | SLCO1B1 | *5 | rs4149056 | ADR - Myopathy | No recommendation |
| Sulfonylurea | CYP2C9 | *2 | rs1799853 | Efficacy and Dosage | No recommendation |
|  |  | *3 | rs1057910 | adjustment |  |
| Thiopurine<br>Methyltransferase | TPMT | c.460G>A, *3B<br>c.238G>C, *2<br>c.719A>G, *3C | rs1800460<br>rs1800462<br>rs1142345 | Toxicity and ADRs | Genetic testing<br>recommended |
| Warfarin | CYP2C9 | *2<br>*3 | rs1799853<br>rs1057910 | Efficacy and Dosage<br>adjustment | Actionable PGx |
|  | VKORC1 |  | rs9923231 |  |  |

<sup>a</sup> PharmGKB evaluated the pharmacogenetic (PGx) information provided by U.S. FDA approved drug labels, and assigned each test with a level of recommendation according to the level of clinical evidence.[6] The levels begin from genetic testing required, genetic test recommended, Actionable PGx to Informative PGx.

The FDA considers that pharmacogenetic tests for five of the 12 drugs provide actionable results (fluorouracil, peginterferon alpha, phenytoin, pseuodocholinesterase deficiency and warfarin); genetic testing is required for one (abacavir); recommended in two (clopidogrel and thiopurine methyltransferase); one provides informative results (proton pump inhibitor) and the remaining three tests have no FDA recommendations (acetaldehyde, simvastatin, sulfonylurea).

For most drugs, 23andMe tests the variants listed by the FDA, but some differences exist. For phenytoin, 23andMe reports on drug sensitivity from *CYP2C9*2* and *\*3* variants, while the U.S. FDA provides pharmacogenetic information on HLA-B*15:02, which confers increased risk of life threatening hypersensivity reactions like Stevens-Johnson syndrome (SJS) and toxic epidermal necrolysis (TEN).[11] CPIC guidelines strongly recommend alternative treatments for patients with HLA-B*1502 positive status,[12] but 23andMe does not test for HLA-B*1502.

### Differences between Ethnicities

Many pharmacogenetic variants have frequencies that vary substantially by ethnicity, and different patterns of LD in associated regions, meaning that the relevance and interpretation of tests is not homogeneous across populations. For example, common genetic polymorphisms in *CYP2C19* and *CYP2C9* alter ability to metabolise drugs.[13] Individuals with *CYP2C19*2, *3, *4, *8* and *CYP2C9*2, *3* polymorphisms are poor metabolisers and may require dose reductions or an alternative drug. The frequency of *CYP2C19* poor metaboliser genotypes is highest in individuals of East Asian ancestry (14%), and is much lower in those of African ancestry (4%) or European ancestry (2%).[14] Conversely, *CYP2C9*2* and *\*3* variants are common in European populations, with frequencies of approximately 14% and 8% respectively. These variants have lower frequency in African and East Asian populations.[12]

Pharmacogenetic testing for abacavir hypersensitivity is well-established, with the drug label stating “All patients should be screened for the HLA-B*57:01 allele prior to initiating therapy”.[15] Two large studies, PREDICT-1 and SHAPE, have assessed the clinical utility of this test (Table 1).[16, 17] The studies confirmed strong association between HLA-B*57:01 carrier status and immunologically confirmed hypersensitivity reactions, with sensitivity of 100% and specificity of >96% in both European and African ancestries. 23andMe reports test results based on SNP rs2395029, which is in strong linkage disequilibrium with HLA-B*57:01. However, rare recombination events occur in some populations[18] and there is incomplete LD between rs2395029 and HLA-B*57:01:[19–21] 23andMe report rs2395029 heterozygotes as having a 94% chance of carrying HLA-B*57:01. In addition, little is known about the predictive value of the SNP-based test in non-European populations. The rs2395029-G allele is extremely rare in the African population (~0%), and is only present in 1% of the East Asian population. LD between HLA-B*57:01 and rs2395029 varies by population, with perfect LD in Han Chinese and Tamil Indians, but lower in Southeast Asian Malays (r^2^=0.75). These SNP test results should therefore be interpreted with caution, particularly in non-European populations.

Peginterferon alpha (PEG-IFN-alpha) and ribavirin (RBV) combined therapy is the standard 48-week treatment for the chronic infection of hepatitis C virus (HCV), but 25-40% of patients fail to respond to these treatments. Genome-wide association studies identified a significant association of rs8099917 with treatment response, with the G allele increasing risk of failure to response. Carriers of the rs8099917 TT genotype have a 2-3 fold increased chance of responding to therapy.[22] A second SNP rs12979860, not reported by 23andMe, may have better predictive value in some populations.[23] 23andMe reports only rs8099917, stating that “because all of these SNPs are so closely linked to each other, they are all probably representing the same effect”.[24] However, the allele frequencies and linkage disequilibrium between these variants differ across populations (Table 2), so while these SNPs are in strong LD in East Asian populations, the LD is much weaker in African populations. The validity of this test may therefore depend on ethnicity of the 23andMe customer, although it is likely that neither SNP is the causal variant.

**Table 2:** Population genetics of SNPs rs8099917 and rs12979860, both associated with response to hepatitis C treatment. Minor allele frequencies (MAF) calculated using 1000 Genomes data; LD parameters calculated using LDlink.[8, 9]

| Population | rs8099917<br>T/G MAF G | rs12979860<br>C/T MAF T | D' | r<sup>2</sup> |
| --- | --- | --- | --- | --- |
| European | 0.17 | 0.31 | 0.97 | 0.43 |
| East Asian | 0.08 | 0.08 | 0.99 | 0.92 |
| African | 0.04 | 0.67 | 1.00 | 0.02 |

Frequencies of the rs9923231 T allele, which increases sensitivity to Warfarin, vary across ethnicities. In 1000 Genomes phase 3 data,[8] the frequency is highest in Europeans (39%), and the T allele is much rarer in South Asians (15%), East Asians (12%), and Africans (5%). For acetaldehyde toxicity, the *ALDH2*2* (rs671, G>A) variant is mainly found in East Asians and is rare in other ancestries.

For 5-FU toxicity, the frequency of *DPYD*2A* is low in all populations. It is almost non-existent in African-American and Japanese populations, and has frequency of 0.91% and 0.47% in Dutch and German populations respectively.[25]

For Simvastatin-induced myopathy, the frequency of the minor allele of rs4149056 is relatively low across all ethnicities, with the highest in Middle Eastern population (5%), followed by Caucasian (1%). It is rarely observed in Asian and African populations.[26]

23andMe genotyping captures four of the five non-functional alleles which reduce TPMT activity. The frequency of the functional *1 allele ranges from 0.925 in Mexicans to 0.983 in Asians^18^ with frequencies of the non-functional markers also varying by ethnicity. The other non-functional alleles are not tested for by 23andme, but have a frequency of < 0.1% in all populations tested.[27]

### Limitations of 23andMe pharmacogenetic tests

For a number of the pharmacogenetic tests performed by 23andMe, the most recent literature has additional information on associations, making 23andMe’s tests incomplete in their assessment of ADRs or efficacy.

For 5-FU, 23andMe tests only for the *2A variant, but as 23andMe state on their website, further variants in the gene are associated with 5-FU toxicity. For example, recent studies have shown that *DPYD*13,* rs67376798, c.1679T>G and c.1236G>A/HapB3 are also associated with 5-FU toxicity.[27-29]

For pseudocholinesterase deficiency, 23andMe provides reports on three variants, fluorine resistant 1 (F1), fluorine resistant 2 (F2) and dibucaine resistant mutation (A). Other mutations in *BCHE* with similar effect are not reported by 23andMe, such as K-variant, silent-1, silent-2 and silent-7.[30] The protocol for selecting the specific variants to be tested by 23andMe is not described and only two citations are listed for this particular drug response, and which does not meet their selection criteria.[3]

For acetaldehyde toxicity, 23andMe reports on rs671, which is mainly present in East Asians. An additional variant rs1229984, A>G in *ADH1B,* increases risk of oesophageal cancer and is present in multi-ethnic populations. Cui R *et al,* 2009 found that the combination of *ALDH2*2* and *ADH1B* variants with smoking and drinking significantly increases the risk of oesophageal cancer by 189-fold.[31]

Assessing variation in TPMT is difficult and 23andMe states that their current technology cannot distinguish between the genetic changes in the same or different copies of the *TPMT* gene. For example, it is not possible to differentiate between one copy of *3A (reduced function) and copies of *3B and *3C on different chromosomes (no function). Although it is rare to carry both *3B and *3C variants, incorrect interpretation of test results could lead to dosing errors.

## Discussion

Adverse events from drug hypersensitivity currently place a large burden on healthcare costs and preventable adverse drug events could cost the UK NHS up to £2.5 billion per year.[32] Some of these events are associated with genetic factors, and so may be preventable through pharmacogenetic testing. DTC pharmacogenetic testing may have a role to play in either raising awareness of pharmacogenetic testing, or providing customers with preliminary information, which could be followed up with healthcare professionals. However, these benefits need to be balanced with the risks involved in providing medical information that is not communicated through a healthcare professional.

The test provided by 23andMe highlight the challenges of pharmacogenetic testing, some of which are unique to a SNP-based genotyping platform used by the DTC company, others of which are applicable to any technology, and any method of delivery. We compared the 12 tests provided by 23andMe with recommendations from the FDA and CPIC, finding several differences between the two sources. 23andMe report on some pharmacogenetic tests for which FDA provided no recommendation (simvastatin, sulfonylurea and acetaldehyde); these tests are supported by research publications and therefore (mostly) meet the selection criteria specified by 23andMe. 23andMe only reports on a subset of tests with FDA recommendations or CPIC guidelines. Some omissions may be due to difficulties in tagging the relevant pharmacogenetic variant. For example, 21% (44/204) of the tests in the FDA Table of Pharmacogenomic Biomarkers in Drug Labelling are for *CYP2D6* alleles. These cannot be assessed fully with a genome-wide genotyping chip due to gene deletions and sequence similarity with pseudogenes, and this may explain the lack of CYP2D6 tests in 23andMe’s portfolio. Some tests provided by 23andMe provide an incomplete assessment of risk variants within the target genes (*DPYD, ADH1B*). For phenytoin, the variant tested by 23andMe differs from recommendations by the FDA. The FDA’s table lists tests for *CYP2C9, CYP2C19* and *HLA-B*\*15:02, with CPIC providing guidelines for joint tests across *CYP2C9* and *HLA-B*\*15:02, while 23andMe reports results on *CYP2C9.* This omission is potentially serious since *HLA-B*\*15:02 carriers are at increased risk of SJS/PTEN, and use of phenytoin in not recommended. Similarly, carbamazepine-induced SJS/PTEN occurs in *HLA-B*\*15:02 carriers and this test is not provided by 23andMe. This pharmacogenetic test is particularly relevant in East Asians where HLA-B*15:02 frequency is highest. Two SNPs in strong LD with HLA-B*15:02 have been identified. A sample of 45 Han Chinese from Beijing, China showed r^2^=1 with rs3909184 and rs2844682,[33] but LD with rs3909184 was much weaker in an Asian Pacific Islander population (sensitivity 32%; no r^2^ value given). These population-level differences in LD between HLA alleles and tagging SNPs make it difficult to implement a pharmacogenetic test using only SNP data from genotyping arrays, particularly in a DTC setting which does not allow for subtleties in test interpretation. This highlights the point that fully comprehensive pharmacogenetic testing would require a combination of SNP genotyping or sequencing, plus HLA serotyping.

For drug metabolising enzymes, 23andMe provide reports on *CYP2C19* for clopidogrel and proton pump inhibitors, but not for other drugs such as the widely prescribed selective serotonin reuptake inhibitor anti-depressants (SSRIs). CYP2C19 tests are listed on the FDA table for SSRIs, and are available in a commercial test by AssureX.[34] Guidelines for *CYP2C19* pharmacogenetic tests are provided by CPIC (8 drugs) and the Dutch Working Group (12 drugs), each of which includes the test provided by 23andMe.

One of the challenges in pharmacogenetics is the diverse allele frequencies across populations. Different allele frequencies may make the test only relevant in a single population, as in testing for phenytoin and HLA-B*15:02 which is mainly found in Asian populations. Differing allele frequencies for the causal variant does not affect interpretation of the pharmacogenetic test result, but different linkage disequilibrium may invalidate interpretation of a tagging SNP, as seen with rs8099917 and Hepatitis-C treatment response. Without further work refining causal SNPs at such loci, it is probable that pharmacogenetic tests will provide different degrees of sensitivity across ethnicities.

We have focused on DTC company, 23andMe, since they are the largest provider in this market, and are commendably open about the exact SNPs genotyped, and the reports generated for each pharmacogenetic test – therefore enabling scrutiny of the service offered. A 2012 paper reviewed the DTC pharmacogenetic testing companies in the market, identifying eight companies that provided at least one pharmacogenetic test.[35] Some of the companies surveyed require tests to be ordered through a clinican (Genelex, Pathway genomics), or are no longer in the market (Navigenics). Geneplanet now provides DTC pharmacogenetics for six drugs or drug classes (Clopidogrel, Metformin, Omeprazole, Perindopril, Statins, Warfarin), and Theranostics for two drugs (Clopidogrel, statins) but neither company gives detailed information on the tests or reported outcomes and so are not included here.

DTC pharmacogenetic testing has the potential to benefit patients by educating individuals about genetics and providing preliminary information that patients could follow up with a medical professional. To 23andme’s credit, it is made very clear on their public website, and in reports for registered customers, that they do not provide a medical service, instructing customers to “not stop, start or change a drug regimen without consulting a doctor”. In addition, 23andMe provides detailed information about the test performed, the evidence for gene-drug association, and the implications of test results. In addition to clinical-related information, full SNP genotypes are returned. Pharmacogenetics is a particularly relevant focus for DTC testing since the test has value throughout life, and need be performed only once. This makes implementation of a pre-emptive pharmacogenetic testing panel potentially cost-effective, since the results can be interrogated when any new drug is prescribed.

The service offered by 23andMe has several important limitations, and for the service offered to be more effective, we would recommend a number of changes:

1. Better mechanisms should be in place to ensure that tests reflect the latest science, to ensure tests do not become outdated. Pharmacogenetic research can move quickly – and producing out-of-date reports raises ethical questions since reports may be invalid based on updated research results.
2. Better consideration and documentation of differences across ancestry. For example, the literature shows substantial differences in sensitivity and specificity of tests for Hepatitis-C treatment response between ancestries. This should be fully reported by 23andMe to ensure all test users understand the ancestry-specific limitations of testing. In the absence of research linking the tested SNP to the response in a given consumer’s ethnicity, this limitation should be communicated clearly.
3. More consideration of what tests should be reported. Many 23andMe customers will not be knowledgeable about the relative importance of the different tests. Why provide results from tests that are uninformative – at least according to FDA – when this could cause unnecessary concern/anxiety for customers?

Sequencing technologies may be required to achieve these aims, as the current genotyping arrays do not always offer the variants required to perform effective testing, and would enable a wider range of important drug metabolising enzyme variation to be detected.[36] In addition, rare variants not identified in the literature will continue to provide challenges to pharmacogenetic testing on a large scale, and reduce the specificity of tests performed.

## Acknowledgements

This paper represents independent research funded by the National Institute for Health Research (NIHR) Biomedical Research Centre at South London and Maudsley NHS Foundation Trust and King’s College London and by the NIHR Biomedical Research Centre at Guy’s and St Thomas’ NHS Foundation Trust and King’s College London. The views expressed are those of the author(s) and not necessarily those of the NHS, the NIHR or the Department of Health.

